# Effects of social position on subsequent courtship and mating activity in an African cichlid

**DOI:** 10.1101/2024.04.26.591340

**Authors:** Anastasia Martashvili, Sara Jedwab, Lakshita Vij, Avraham Zion Kuighadush, S.G. Alvarado

## Abstract

Within social hierarchies, rank can be dynamic and modulated by changes in molecular and/or physiological substrates. Here, we sought to better understand how social environment and rank shape male spawning behaviors and outcomes in African cichlid fish *Astatotilapia burtoni*. First, using a social dyad paradigm, we generated territorial (T)/ Non-territorial (NT) male pairs. After establishing a stable social hierarchy, the behaviors of the Ts and NTs were recorded and scored. Afterward, pairs were separated and individually moved into a spawning phase, which consisted of a new tank with novel females and no other males where their behaviors were scored. While previous studies have shown how territorial and non-territorial males have unique behavioral profiles, we sought to deepen this interpretation with a focus on the latency of decision-making, and on transition matrices representing enriched sequences of behavior. We found that while the courtship behaviors are shared between stably territorial and ascending males in the spawning phase, only the animals that were territorial in the dyad phase were the ones that were reproductively successful in the subsequent 16 hour spawning phase.

## Introduction

Social hierarchies are prevalent across the animal kingdom and allow individuals within a community to secure resources and mating opportunities. For example, social rank can bestow reproductive potential on an individual, which can lead to variation in behavior, physiology, and gross anatomy. These systems can be developmentally biased or plastic (Laland et al., 2015). For instance, the developmental constraints on reproduction in hymenopterans create a developmental bias that physiologically generates workers who cannot produce fertile offspring in the presence of a queen (Khila & Abouheif, 2008). Similarly, subsequent pregnancies in naked mole rats can lead to vertebrae elongation, leading to larger litters and queen caste determination (Dengler□Crish & Catania, 2009; Thorley et al., 2018). In contrast, social systems that are plastic can shift rank among conspecifics via behavioral transitions. For example, changes in behavior and serotonergic tone or gonadotropin function can shift ranks among *Anolis* lizards (Summers et al., 2005) and African cichlids (Maruska & Fernald, 2018), respectively. While changes in rank can present social opportunities, little is known about how transitions in rank can affect decision-making and complex behavior within a population. For example, how does previous social life history inform changes in a novel social environment, and does a continuum exist between developmentally plastic and biased behaviors?

Here, we studied the African cichlid fish *Astatotilapia burtoni* (Günther 1894), which is endemic to Lake Tanganyika. The species was selected because of its robust social hierarchy, and plasticity in its behavior and morphology in response to its environment. *A. burtoni* has been recognized as an emerging model for the study of social behavior, which has been associated with the diversification of cichlids in East African Great Rift Lakes.

Social status within lek-like communities of *A. burtoni* is linked to male-specific coloration and social behaviors, and is believed to be a consequence of intense sexual selection and male-male competition (Seehausen & Schluter, 2004). In the wild, *A. burtoni* has adopted a lek-like social system, wherein two distinct types of males are observed: territorial (T) and non-territorial (NT) (Fernald & Hirata, 1977), and females visit leks for mating. T males tend to exhibit aggressive defense of specific areas within the habitat, typically against intruding males, whereas NT males refrain from establishing or defending specific territories. T males are commonly larger and display more vibrant coloration than their NT counterparts. Furthermore, these hierarchies and male stereotype behaviors can be induced and replicated in semi-natural environments in the laboratory (Fernald, 1977).

In the laboratory, the behavioral dynamics of *A. burtoni* males have been extensively studied in relation to the neurobiology of social behavior (Maruska & Fernald, 2018). Male *A. burtoni* naturally assume territorial status within their environment (Fernald, 1980), even in the absence of competitors, suggesting that their territorial behavioral displays suppress non-territorial males from ascending in rank. Changes in rank via a social ascent or descent result in rapid shifts in behavior within minutes (Maruska et al., 2013; Fialkowski et al., 2021). For example, the removal of a dominant male from a tank with suppressed NT males presents the social opportunity to ascend to territorial status (Maruska, 2015). Similarly, T→NT transitions are linked to directional changes in behavior that are accompanied by rapid changes in cortisol and immediate early gene expression in the brain (Maruska et al., 2013). Additionally, changes to the physical environment can prime transitions in social ascent/descent via the modulation of growth rates (Hofmann et al., 1999) or morphological coloration (Fang et al., 2022) and their accompanying behavioral profiles (Korzan et al., 2008). Thus, the physical environment can have profound effects on their social environment.

Given the broad developmental plasticity seen in *A. burtoni*, we used the system to ask the question: how do stable and transitioning ranks shape the reproductive behaviors of males? We present a study of T/NT males maintained by a stable dyad paradigm and subsequent exposure to a novel social environment where a focal male can mate with multiple females. Briefly, T/NT male pairs were generated over a 1-month dyad and put into a larger spawning paradigm consisting of a larger tank with ten females where the behavior of the focal male was scored in the first hour. We sought to investigate whether their past experiences in a stable dyad T/NT male would affect their behavioral patterns as territorial individuals in a novel social environment (spawning phase). In addition to examining the incidence of different behaviors, we complement our understanding of these behavioral profiles with analyses of the latency to make decisions and behavioral sequence transitions to identify nuanced differences between behavioral profiles. We anticipate a deeper understanding of these processes will inform our understanding of male social behaviors, how behaviors shape their fitness, and the evolution of behavior in cichlids.

## Methods

### Animals and animal husbandry

Subjects were laboratory-bred cichlid fish *A. burtoni*, derived from stock wild-caught in Northern Lake Tanganyika, Africa (Fernald & Hirata, 1977). All animals in this experiment were bred and reared in tanks that were set up to mimic the natural conditions of Lake Tanganyika. Water was maintained at pH 8, 25°C, salt 280 ppm, while a flow down system enabled a water change twice a day, and the lighting was on a 12-hour day/night cycle (Alvarado et al., 2015a). Fish were fed once a day at 9 AM and kept in tanks lined with brown gravel. At the time of the experiment, all animals were six months to one year old and reared in stock tanks (∼40 males and females in 115 L tanks), and all females were adults capable of spawning as they all had at least one brood previously.

### Experimental setup and behavioral recording

In the social dyad phase of the experiment, two males were age- and size-matched (±2.5-3 mm) and housed with three females for one month (N=10 replicate T/NT pairs) (Fig.1). We controlled for the males’ social life history by choosing males that were suppressed (muted colors) in a stock tank in the presence of a much larger dominant male. Males in the dyad paradigm typically establish status within the first 24 hours and develop behavioral and physical phenotypes consistent with their social status (Maruska, 2015). The dyad tank had a natural substrate comprising brown gravel and half of a terracotta pot functioning as a territory (Fernald, 1977). Social status of each male was determined by looking at pot ownership, body coloration and behavioral displays where the male that occupied the pot, exhibited more vibrant coloration and more aggressive behavior repertoire was classified as territorial (Maruska, 2013). All dyad assays were stable as we observed no rank reversals. At the end of one month, dyad tanks were filmed for one hour with cameras (GoPro Hero8 Black) mounted on tripods (Amazon Basics WT3540). In the second phase of the experiment, directly after the dyad filming concluded, each male was removed from the dyad tanks and put separately into the spawning phase: a new tank with brown gravel, half of a terracotta pot, and ten adult females the focal male had never interacted with before. Filming started immediately, and they were filmed for one hour. Following 16 hours after the filming, mouthbrooding females were counted in each tank. Spawning trials were staggered, with subsequent spawning phases taking place with alternating males of different ranks, which was specifically designed to make sure the females were exposed to both types of males.

**Fig. 1.**
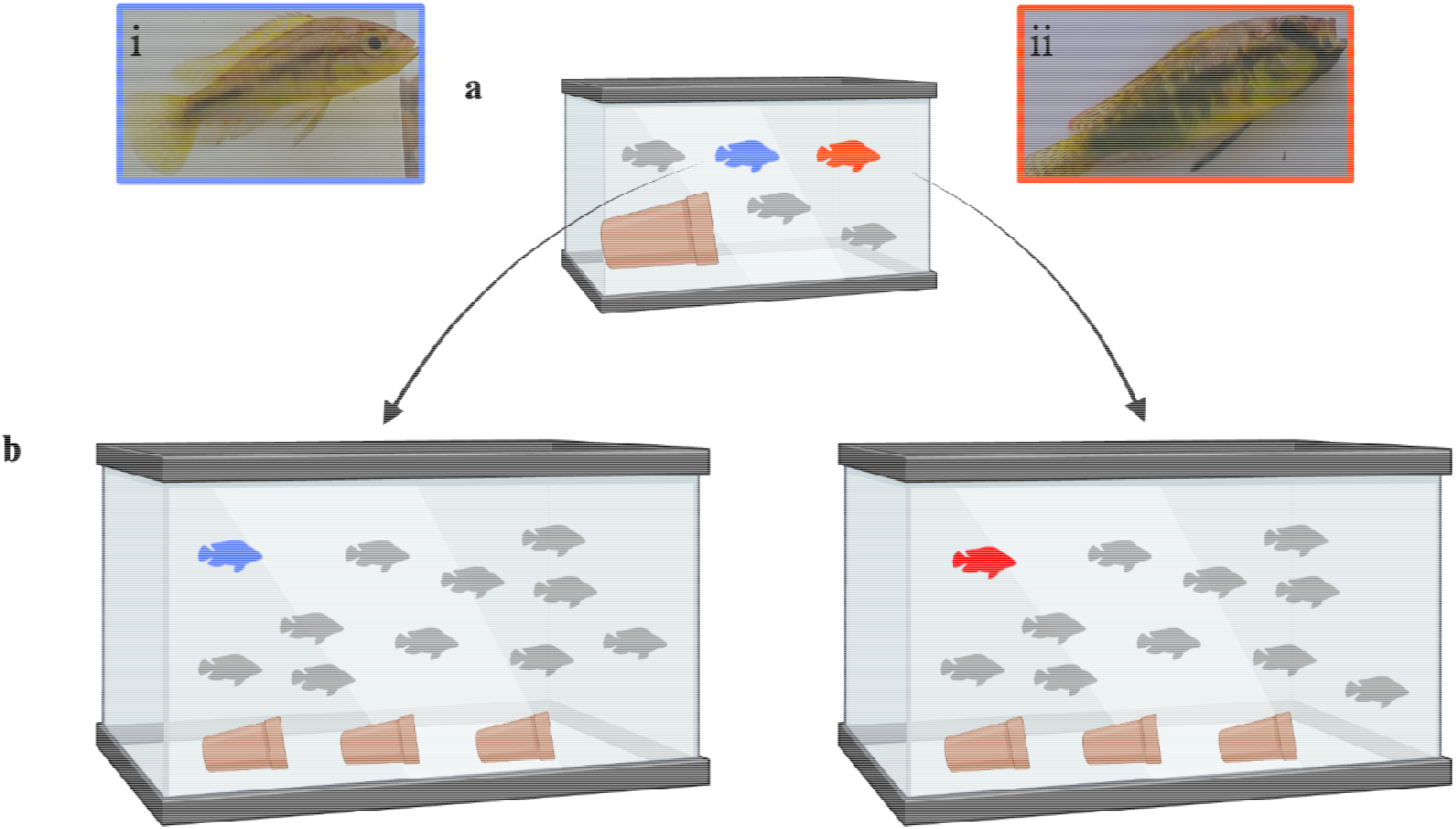
Illustration of the behavioral assays used. a) In dyad assay, two males and three females (gray) were placed in a tank with brown gravel and half of a terracotta pot as a territory (n=10). After a month of induction, males either ascended to NT status (i) or T status (ii) and were moved into a spawning assay. b) in spawning assay, each of the males were put in a tank with three terracotta pots, brown gravel and 10 adult females

### Behavioral Scoring and Analysis

Videos were scored in BORIS event-logging scoring software (Friard & Gamba, 2016) by a blinded scorer using an ethogram previously described (Fernald & Hirata, 1977). Following behaviors were identified: *“attack female”, “chase female”, “dig”, “pot entry”, “pot exit”, “lead swim”, “flee from male”, “flee from female”, “attack male”, “chase male”, “quiver at female”, “quiver at male” and “lateral display”*. Score logs were exported from BORIS in a CSV format. Data wrangling, statistical analyses, and boxplot generation were performed in R (version 4.2.2, 2022-10-31) within RStudio, using the following packages: data.table (version 1.14.8), dplyr (version 1.1.3), ggplot2 (version 3.4.4), ggpubr (version 0.6.0), pwr (version 1.3.0), readr (version 2.1.4), reshape2 (version 1.4.4), stats (version 4.2.2), stringr (version 1.5.0), and tidyverse (version 2.0.0) (R Core Team, 2022; Dowle & Srinivasan, 2023; Wickham et al., 2023a, 2023b; Wickham, 2016; Kassambara, 2023; Champely, 2020; Wickham, 2007; Wickham, 2022; Wickham et al., 2019). After combining dyad and spawn data, to account for the possibility of some individuals being more active than others, relative frequencies were calculated, normalizing behaviors within each subject. Then, paired t-tests were conducted on individual behaviors and behavioral categories, comparing the dyad T with dyad NT and spawn T with spawn NT, respectively. The Benjamini-Hochberg correction, also known as false discovery rate (FDR) correction, was utilized to adjust for the increased likelihood of false positives when conducting multiple hypothesis tests simultaneously, using the p.adjust() function in R (James et al., 2023). For the latency graphs, we created a data frame using base R that reflected the behavior pair and the time taken to transition from one behavior to another, mean, and standard error of the mean (SEM) of each behavior done by each subject in each phase. Python version 3.11.5 and its libraries such as NumPy, pandas, NetworkX, and Matplotlib were used to generate Markov chains to visualize behavioral transition matrices (Foundation, 2023) where nodes represent each behavior, along with their relative sizes reflecting how frequently each behavior occurred.

All figures were finalized using Biorender.

## Results

### Territorial males hold rank by carrying out suppressive agonistic behaviors on non territorial males

In the dyad phase, compared to NTs, Ts undertook more pot-related behaviors such as *“pot entry”* (p<0.01) and *“pot exit”* (p<0.01), reproductive behaviors such as “*chase female*” (p<0.05) and “*quiver at female*” (p<0.05) and aggressive behaviors of “*attack male*” (p<0.01) and ”*chase male*” (p<0.05) (Fig. 2a, Table 1). In contrast, NTs did more “*flee from female*” (p<0.05) and “*flee from male*” (p<0.01) than Ts. When considering incidence of aversive, reproductive, and aggressive categories, Ts did more aggressive ( p<0.01) and more reproductive (p<0.05) behaviors than NTs, and NTs engaged in more aversive behaviors (p<0.01) than Ts (Fig. 2c, Table 2).

**Fig. 2.**
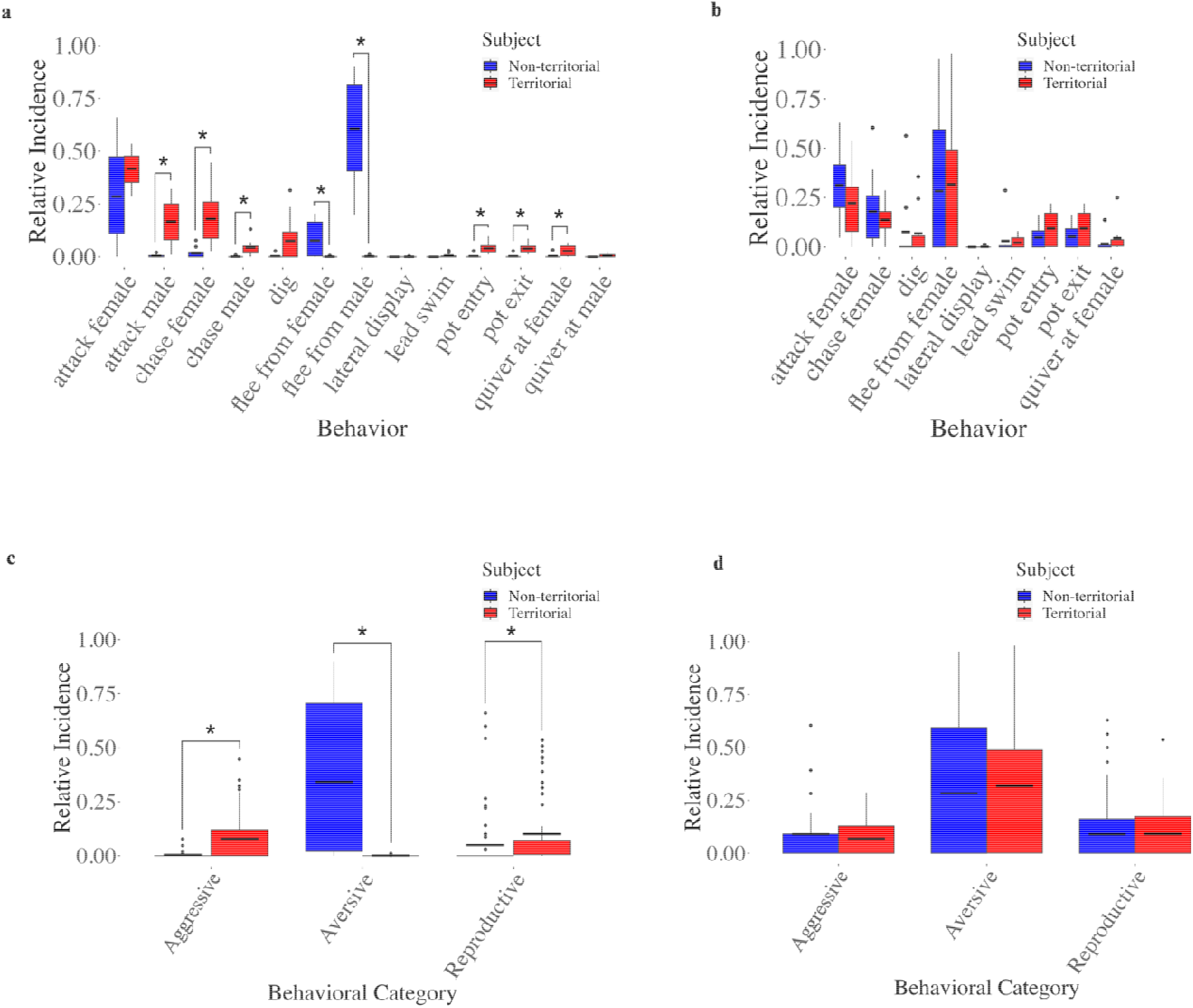
Illustration of individual and categorical behavioral incidences of territorial and non-territorial males in dyad and spawn assays. a) We observed differences between the territorial and non-territorial males in the dyad assay in the following behaviors: “*attack male*” (p<0.01),“*chase female*” (p<0.05),”*chase male*” (p<0.05), “*flee from male*” (p<0.01), “*flee from female*” (p< 0.05), “*pot entry*” (p<0.01), “*pot exit*” (p<0.01),”*quiver at female*” (p<0.05). b) There were no differences in behavior between territorial and previously non-territorial males in the spawn assay c) In the dyad assay, T males engaged in more Aggressive (p<0.01) and Reproductive (p<0.01) behaviors than NTs and NTs engaged in more Aversive (p<0.05) behaviors than Ts. d) Ts and previously-NTs did not differ in the incidence of behavioral categories in spawn assay. The box plot illustrates the distribution of data as follows: the upper edge of the box represents the 75th percentile, the horizontal line inside the box marks the mean, and the lower edge of the box represents the 25th percentile. Additionally, the line extending above the box depicts the range of values from the 75th percentile to the maximum, while the line extending below the box shows the range of values from the minimum to the 25th percentile. Asterisks denote significance (p<0.05)

**Table 1.** Summary of behavioral comparison between territorial and non-territorial subjects in the dyad assay. The table includes the list of behaviors, test statistics of the paired t-test, degrees of freedom, original p values and the Benjamini-Hochberg adjusted p-values. Missing values for a behavior indicate that t test was not conducted due to insufficient data. Additionally, mean and the standard error of the mean (SEM) values are listed for both territorial and non-territorial males. All Statistical analyses were done using relative incidence values

| Behavior | Test statistic | DF | p value | Adjusted p value | MeanT ± SEM | MeanNT ± SEM |
| --- | --- | --- | --- | --- | --- | --- |
| "attack female" | -1.655278 | 9 | 1.32E-01 | 0.143273456 | 0.417 ± 0.028 | 0.316 ± 0.075 |
| "attack male" | -4.775522 | 9 | 1.01E-03 | 0.006549134 | 0.167 ± 0.034 | 0.014 ± 0.004 |
| "chase female" | -3.746706 | 9 | 4.58E-03 | 0.011902481 | 0.179 ± 0.044 | 0.039 ± 0.015 |
| "chase male" | -3.545722 | 9 | 6.26E-03 | 0.013558377 | 0.049 ± 0.012 | 0.008 |
| "dig" | -1.952616 | 9 | 8.26E-02 | 0.107404427 | 0.150 ± 0.056 | 0.027 |
| "flee from female" | 2.676375 | 9 | 2.54E-02 | 0.041203812 | 0.006 | 0.110 ± 0.033 |
| "flee from male" | 7.527429 | 9 | 3.59E-05 | 0.000466456 | 0.006 ± 0.002 | 0.607 ± 0.081 |
| "lateral display" |  |  |  |  | 0.003 | 0 |
| "lead swim" | -1.65806 | 9 | 1.32E-01 | 0.143273456 | 0.018 ± 0.007 | 0 |
| "pot entry" | -4.070404 | 9 | 2.80E-03 | 0.009093336 | 0.040 ± 0.008 | 0.027 |
| "pot exit" | -4.307971 | 9 | 1.97E-03 | 0.008525959 | 0.039 ± 0.007 | 0.027 |
| "quiver at female" | -3.276597 | 9 | 9.58E-03 | 0.017793661 | 0.029 ± 0.009 | 0.032 |
| "quiver at male" | -2.126778 | 9 | 6.23E-02 | 0.090057026 | 0.013 ± 0.003 | 0 |

**Table 2.** Summary of categorical comparison of behaviors between territorial and non-territorial subjects in the dyad assay. The table includes a list of behavioral categories, test statistics of the paired t-test, degrees of freedom, original p values and the Benjamini-Hochberg adjusted p-values. Additionally, mean and the standard error of the mean (SEM) values are listed for both territorial and non-territorial males. All Statistical analyses were done using relative incidence values

| Category | Test statistic | DF | p value | Adjusted p value | MeanT ± SEM | MeanNT ± SEM |
| --- | --- | --- | --- | --- | --- | --- |
| Agressive | -4.809064 | 49 | 1.48E-05 | 4.45E-05 | 0.079 ± 0.016 | 0.004 ± 0.002 |
| Aversive | 4.614858 | 19 | 1.89E-04 | 2.84E-04 | 0.001 ± 0.001 | 0.342 ± 0.074 |
| Reproductive | -2.245977 | 59 | 2.85E-02 | 2.85E-02 | 0.100 ± 0.020 | 0.049 ± 0.018 |

**Table 3.** Results of the paired t-tests comparing behavior transition latencies of dyad T subjects to spawn T subjects. The table includes the test statistics of the paired t-tests, degrees of freedom, original p values and the Benjamini-Hochberg adjusted p-values. Additionally, mean and the standard error of the mean (SEM) values are listed for both dyad and spawn territorial males. Behavioral transitions that did not have sufficient data to run t-tests are not listed. All Statistical analyses were done using relative incidence values

| Transition | Test statistic | DF | p value | Adjusted p value | MeanDyad ± SEM | MeanSpawn ± SEM |
| --- | --- | --- | --- | --- | --- | --- |
| "attack female"\|"attack female" | -0.96972903 | 436.54087 | 0.332718543 | 0.44917003 | 10.941±0.9283 | 12.296±1.0444 |
| "attack female"\|"pot entry" | 2.04122222 | 54.618445 | 0.04607236 | 0.21406929 | 13.9429±2.3474 | 8.5636±1.1976 |
| "pot entry"\|"pot exit" | 1.93510836 | 111.95959 | 0.055499446 | 0.21406929 | 10.1249±1.5072 | 6.7515±0.8759 |
| "pot exit"\|"attack female" | 0.58927945 | 32.573021 | 0.559739058 | 0.65708498 | 10.0526±5.1015 | 6.928±1.446 |
| "pot exit"\|"dig" | -1.04133892 | 8.931563 | 0.325084565 | 0.44917003 | 2.2308±0.7438 | 4±1.5275 |
| "pot exit"\|"pot entry" | -3.0280994 | 18.96415 | 0.006927758 | 0.09352473 | 2.625±0.8954 | 19.2897±5.43 |
| "chase female"\|"attack female" | -2.57139223 | 60.301146 | 0.012612327 | 0.11351095 | 10.0084±1.5087 | 21.2216±4.0915 |
| "attack female"\|"chase female" | -1.99499366 | 68.858439 | 0.050002784 | 0.21406929 | 9.2113±1.56 | 16.7599±3.4472 |
| "chase female"\|"chase female" | -3.75116513 | 45.177493 | 0.000499398 | 0.01348373 | 7.247±0.9677 | 26.3764±5.0069 |
| "lead swim"\|"attack female" | -1.6415558 | 8.439885 | 0.137337555 | 0.33710127 | 3.25±0.8399 | 11.6667±5.058 |
| "dig"\|"pot entry" | -1.45418168 | 8.779604 | 0.180691676 | 0.38113315 | 3.1111±0.6111 | 5.75±1.7087 |
| "dig"\|"dig" | -2.50423315 | 35.726022 | 0.016978963 | 0.114608 | 3.1382±0.3452 | 8.6571±2.1766 |
| "dig"\|"attack female" | -1.3964166 | 12.219782 | 0.187437612 | 0.38113315 | 5.7±0.8485 | 18.1538±8.878 |
| "chase female"\|"pot entry" | 0.66576666 | 43.385106 | 0.50908474 | 0.62478582 | 18.1704±5.1766 | 13.8284±3.9669 |
| "dig"\|"chase female" | -1.05199768 | 6.643201 | 0.329554073 | 0.44917003 | 5±0.7303 | 8.4286±3.1762 |
| "quiver at female"\|"attack female" | -0.01908912 | 51.930446 | 0.984843164 | 0.98484316 | 9.2889±3.3716 | 9.3804±3.4089 |
| "attack female"\|"dig" | -1.28602901 | 27.404858 | 0.209193018 | 0.38113315 | 7.2973±0.7562 | 9.3333±1.3909 |
| "chase female"\|"dig" | 0.82204619 | 5.323865 | 0.446292399 | 0.57380451 | 8.5385±1.4215 | 6.25±2.3936 |
| "attack female"\|"lead swim" | -2.13766701 | 4.957109 | 0.086046551 | 0.26050741 | 2.3333±0.8819 | 5.5±1.1902 |
| "pot exit"\|"chase female" | -0.12834622 | 34.648488 | 0.898617077 | 0.97050644 | 5.7165±2.6893 | 6.1817±2.4303 |
| "attack female"\|"quiver at female" | -1.73105322 | 27.338266 | 0.094715646 | 0.26050741 | 5.79±1.0895 | 14.6511±5.0016 |
| "chase female"\|"quiver at female" | 0.37147081 | 17.309542 | 0.714794794 | 0.80414414 | 13.2489±5.2048 | 10.7385±4.3107 |
| "quiver at female"\|"chase female" | 0.06158099 | 9.415628 | 0.952183982 | 0.98484316 | 10.2495±7.1063 | 9.7764±2.9181 |
| "chase female"\|"lead swim" | 2.35277811 | 3.128147 | 0.096484228 | 0.26050741 | 13±4.4159 | 2.5±0.6455 |
| "quiver at female"\|"quiver at female" | -1.32903174 | 10.597726 | 0.21174064 | 0.38113315 | 5.4147±1.9377 | 9.7±2.5773 |
| "pot entry"\|"attack female" | 1.16925544 | 4.084463 | 0.305978193 | 0.44917003 | 23.8192±15.6882 | 5.3775±1.6255 |
| "attack female"\|"pot exit" | 1.51856261 | 2 | 0.268197218 | 0.44917003 | 13.1943±8.6887 | 0 |

### T/NT rank behaviors in a spawning assay are similar despite robust differences in a dyad

Qualitatively, T and NT behaviors were discretely different in the dyad but very similar in the spawning phase (Fig. 3). This was illustrated with T males engaging in agonistic and reproductive interactions among conspecifics (Fig. 3a), whereas nearly all NT behaviors transitioned to flee (Fig.3b). In contrast, in spawn assay, both T and NT-ranked males illustrated the same behavior patterns (Fig. 3b and d). As we calculated the difference between the probability values of each behavioral transition for T and NT subjects in the dyad assay, we screened the results with a 10% threshold, after which, we took into account how frequently the subjects engaged in each of the behaviors within transitions which also had 10% threshold (see Supplemental Tables 2 and 3). All reported results in this section are those that fell above the threshold. In the dyad phase, the following behaviors were most commonly performed by the Ts: “*attack female*”, “*attack male*”, “*chase female*” and “*dig*”. Ts were 59% more likely than NTs to follow up “*dig*” with “*dig*”, 37% more likely than NTs to first “*chase female*” and then “*chase female*” again, 14% more likely than NTs to follow up “*attack male*” with “*attack male*” again, 10% more likely than NTs to “*dig*” and then “*attack female*”, 14% more likely than NTs to “*attack femal*e” and then “*chase female*” and 14% more likely than NTs to “*attack male*” and then move to “*chase male*”. In contrast, the most frequent behaviors that NTs performed were “*flee from male*” and “*attack female*”. NTs were 40% more likely than Ts to “*flee from male*” and then “*flee from male*” again, 33% more likely than Ts to “*attack female*” and then “*flee from male*” and 20% more likely than Ts to transition from “*flee from male*” to “*attack female*”.

**Fig. 3.**
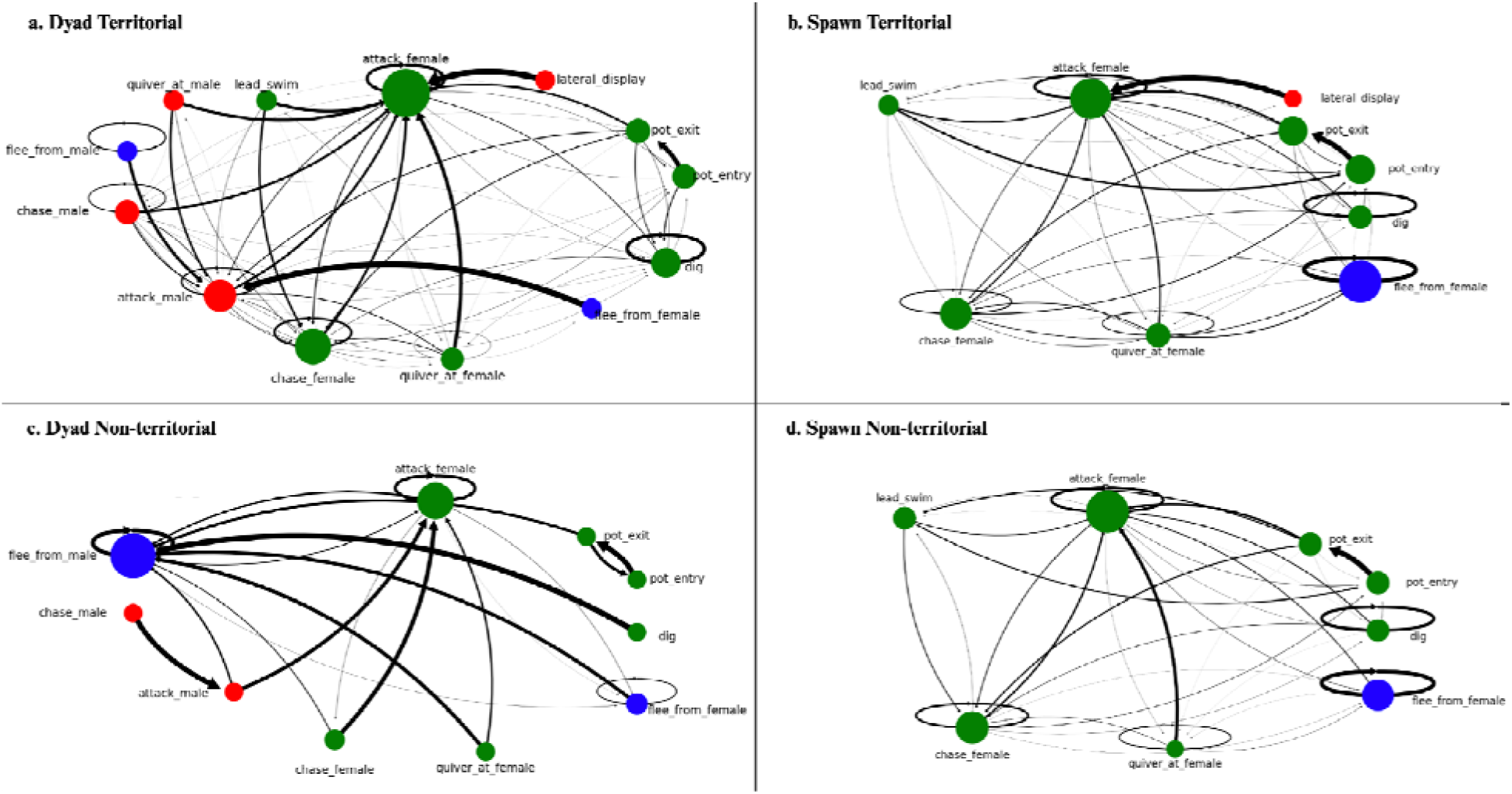
Illustration of the Markov chain analysis conducted to examine the transition probabilities of a) Ts in dyad assay, b) Ts in spawn assay, c) NTs in dyad assay and d) NTs in spawn assay. See Supplemental Figure 3 for a sample Markov chain diagram that clarifies each component, providing useful context for the more detailed figure. There are 9 nodes, each representing a behavior. The color of the node corresponds to the category of the particular behavior: aggressive in red, aversive in blue and reproductive in green (for ethogram, see Supplemental Table 1). The relative sizes of the nodes reflect how frequently each behavior occurred. In any given transition, the line originates from the starting behavior and ends at the follow-up behavior with an arrow. The thickness and amount of shading of the connecting lines between nodes represent how likely one behavior is to succeed another: the thicker and darker the line, higher the probability. If a certain node is missing a connecting line to a node, that means the probability of that particular behavioral transition is 0. Overall, the analysis reveals a drastic change in behavioral transitions of previously-NTs once they were allowed to ascend

We identified differences in behavioral sequences and transitional probabilities between the Ts and NTs in the subsequent spawning phase (Fig.3b,d). As with the previous assay, we studied differences in transitional probabilities between T and previously NT males with the same parameters and found four behavioral transitions in total fell over the 10% threshold consideration (see Supplemental Tables 4 and 5). Territorial males were 13% more likely than previously NTs to “*chase female*” and then “*flee from female*” and 11% more likely than previously NTs to follow up “*pot exit*” with “*pot entry*”. In contrast, previously NTs were 15% more likely than Ts to “*flee from female*” and then “*chase female*” and 14% more likely than Ts to follow up “*chase female*” with “*chase female*” again.

In total, 12 behavioral transitions were performed only by the territorial males in the spawning assay (see Supplemental Table 6). Of those, only one behavioral transition - “*lateral display*” to “*attack female*”-had a probability value that resulted in a difference of over 10%, however, “*lateral display*” was only done by Ts less than 1% of the time. One behavioral transition was performed only by the previously NTs, however, neither the probability nor the incidence values were over 10%. Ts were more likely to transition between more combinations of behaviors than NTs, however, despite some differences, the behavior repertoires of Ts and previously NTs were similar. Notably, there was a drastic difference between the transitional probabilities and behavioral repertoire between the dyad and spawn phase NT subjects. NTs went from barely engaging in any behaviors or transitions to almost the same behavioral repertoire as the T males in the spawning paradigm.

### When presented with spawning opportunities territorial and non territorial males both show similar latencies in decision making

In the spawning phase, there was no statistical difference in the latency of any behavior initiations between T and previously NT males (see Supplemental Figure 1). Of note, NTs did not perform *“lateral display”*, while Ts did so twice. Furthermore, the time taken to transition between two behaviors by Ts and NTs was analyzed. Dyad Ts exhibited the same decision making latencies as dyad NTs and spawn Ts exhibited same decision making latencies as spawn NT subjects (see Supplemental Figures 2a,2b). However, when comparing the decision-making time of the same subjects across dyad and spawning phases, we found that Ts, when they were in dyad phase, were much faster transitioning from “*chase female*” to “*chase female*” compared to when they were in spawn phase (p<0.05) (see Supplemental Figure 2c, Table 3). NTs exhibited no differences in transition times between behaviors when moved from dyad into spawning assay (see Supplemental Figure 2d). It is important to note that in the dyad phase, NTs engaged in fewer behaviors in general, therefore, there were behavioral transitions that occurred at very low incidence and that data was insufficient for the paired-t test analysis. Most of the territorial males reproduced (1-2 broods), however, none of the previously non-territorial males were successful in reproduction.

## Discussion

This study presents territorial and non-territorial behavior sequences within a dyad and follows a change into a novel social environment in the form of a spawning paradigm. While the courtship behaviors are shared between these two groups in the spawning phase, only the territorial animals in the dyad phase succeeded in being reproductive in the subsequent spawn phase. When presented with a social opportunity, behaviors associated with territoriality in ascending NT males were reinstated, provided the social opportunity consistent with previous reports (Maruska & Fernald, 2010). Our findings remain consistent with the behaviors that occur during a social ascent in *A. burtoni*, along with the inclusion of decision-making latencies and behavioral sequences that illustrate the similarities between T and ascending NT males. In the spawning phase, T and NT transitional probabilities were similar.

However, Ts exhibited more complexity in transitions, which could be the result of the novelty of the environment and females (Fig.3), which may be analogous to social novelty behaviors reported in other teleost systems such as zebrafish (Barba□Escobedo & Gould, 2012; Nascimento & Maximino, 2023). In zebrafish, social behaviors such as shoaling are typically enriched among novel conspecifics. In *A. burtoni*, we suspect that the same phenomenon is modulated via a broader complexity of behavioral interactions. This complexity can be illustrated by social network centrality differences seen between T and NT males within a population where T males suppress NT males and their behaviors (Rodriguez-Santiago et al., 2020). We suspect our findings could translate to behavioral patterns between males in the wild. *A. burtoni* are lekking fish, often establishing several adjacent territories that shoaling females visit for mating opportunities in diverse visual ecologies. Many behavioral patterns among T males appear to play a suppressive role that delays the ascent of NT males. However, male behavior in the absence of a suppressor (NT spawning assays) show similar behavioral patterns with T males.

None of the NT males sired any broods suggesting that female preference for dominant-looking males may be shaping the outcomes of these experiments. We suspect that female preference may prime subordinate ranked male opportunity during ascension, eventually leading to the pigmentation changes females are attracted to or other nonvisual cues that can take up to a month to occur. For example, territorial males typically show conspicuous coloration with bright blue or yellow body coloration, a red humeral blotch behind the operculum, a black eyebar, and high contrasting stripes whereas non-territorial males show none of these colors, looking more cryptic like females (Maruska & Fernald, 2018). Since T/NT males were rotated between spawning tanks to control for the exposure of ranked males to females, females had the opportunity to see what a T male looks like (compared to an NT), shaping their overall preference for dominant males and their colors. We suspect that this suggests that plasticity animal coloration (known to occur in *A. burtoni* (Fang et al., 2022) in the patterns tied to territorial and non-territorial rank (Peroš et al., 2024) may be important for female preferences (Dijkstra et al., 2024). This is partially reinforced by data where females show avoidance and agonistic interactions with male androgen receptor mutants that show reduced body coloration (Howard et al., 2024).To females, such colors are perceived by a visual system tuned to their reproductive state (Butler & Maruska, 2021), possibly affecting ascending NT reproductive success as it develops these more “attractive” color patterns.

It is important to note that transition matrix analysis, like other probability measures, has a shortfall in that it only accounts for incidence and not duration. Intuitively, it is easy to consider that subjects spend more time performing behaviors that they are more likely to engage in, but additional behavioral experiments need to be conducted to confirm this. Previous studies have found that ascending males exhibited different patterns of aggressive behavior compared to stable T males (Maruska & Fernald, 2010; Alward et al., 2019). Specifically, ascending males were less likely to follow aggression with courtship compared to territorial males. However, this distinction diminished when longer intervals between actions were considered, suggesting that ascending males can adopt typical territorial behaviors, but with slower transitions from aggression toward courtship. In our study, however, there was no difference in the transition latency of any behaviors between T and previously-NT males (See Supplemental Figure 2).

Behaviors involved in the highest number of combinations for each subject may suggest the subject’s tentative motivations. For Ts in the spawning assay, *“attack female”* is involved in the highest number of combinations (16) and 26% of all behaviors, suggesting that affiliative interactions with females indicate either social suppression and/or mating (see Supplemental Table 7). In studies examining the behavioral sequences underlying courtship, mating behaviors are more interconnected involving a series of leading, quivering, and pecks at a spawning site, which was not observed here (Juntti et al., 2016). This suggests that the initial repertoire of behaviors had focal males exploring their novel ambient social environment, with subsequent spawning happening after filming in T males. Similarly, behavior central for the NTs in the spawning assay seems to also be *“attack female”* (13 combinations), comprising 34% of relative behaviors. After the central behavior, the most common behaviors for Ts are “*flee from female*” (28%), *“pot entry”* (11%), *“pot exit”* (10%), and “*chase female*” (13%), and for NTs are “*flee from female*” (17%) and *“chase female”* (13%).

Consistent with other reports, our data shows that territorial males tend to exhibit higher reproductive success, which has been attributed in part to elevated levels of testosterone and 11-ketotestosterone compared to non-territorial counterparts (Maruska et al., 2011; Maan & Sefc, 2013; Maruska, 2015). This is true for other cichlid species such as *Neolamprologus pulcher*, where dominant males are breeders and subordinate males are helpers and the males, as they ascend to being a breeder, exhibit higher androgen levels as well as gonadal investment (Fitzpatrick et al., 2008). Similarly, female preference in *A. burtoni* is largely shaped by male reproductive physiology, with secreted androgens playing an important role in female affiliation and mating bouts (Clement et al., 2005; Kidd et al., 2013). In contrast, none of our ascending NT males showed any reproductive success (evidenced by a tally of mouthbrooding females from spawning tanks). Given that both NT and T males have similar sperm velocities (Kustan et al., 2012) and robust behaviors related to mating, it is difficult to ascertain why NT males failed to reproduce. Since hormonal changes in 11-KT in ascending males occur most robustly after 72 hours (Maruska & Fernald, 2010; Butler & Maruska, 2021), the window for reproduction would have occurred after the animals were sacrificed. Additionally, female-directed behaviors toward the focal male may have also shaped their reproductive success.

## Conclusion

In this study, we examined the patterns and the impact of prior social status on the behavior of stably territorial and ascending male subjects. Previous research has identified distinct behavioral profiles based on male status; here, we aimed to expand on these findings by examining decision-making latency and employing transition matrices to detail sequences of behavior. Ts and ascending males employed very similar behavioral profiles in the spawning phase. Furthermore, our results indicate that while courtship behaviors occurred with equal incidence for both stably territorial and ascending males during the spawning phase, only those males that had maintained territories during the dyad phase were successful in reproducing in the subsequent spawning phase. Future research should focus on documenting and analyzing female behavioral patterns to assess whether they interact differently with males of different social pasts. Overall, the results of this study suggest that previous social experiences can shape reproductive outcomes in different environments while not affecting male-specific stereotypical social behavior.

## Acknowledgments

We would like to thank funds provided by the Queens College Foundation, National Science Foundation (Award # 1921773), and The Professional Staff Congress of the City University of New York (TRADA-50-228) for funding provided to SGA, LV, and SJ and The Cognitive Neuroscience Master’s program at the Graduate Center which supported AM. Special thanks to Andrew Claros who assisted with the development of these tools for behavioral analysis. Additional thanks to Dr. Maral Tajerian for revising this manuscript and providing constructive feedback on its writing.

