## Supplemental Information for "Effects of social position on subsequent courtship and mating activity in an African cichlid"

Hydrobiologia: Advances in Cichlid Research VI: Behavior, Ecology and Evolutionary Biology

Anastasia Martashvili^1^, Sara Jedwab^2^, Lakshita Vij^2^, Avraham Zion Kuighadush^2^, Alvarado S.G.^1,2^

1. **Cognitive Neuroscience Program, Graduate Center of City University of New York, New York, NY 10016**
2. **Department of Biology, Queens College City University of New York, 65-30 Kissena Blvd, Flushing, NY 11361, USA**

| Behavior | Category | Description |
| --- | --- | --- |
| *“attack female”* | Reproductive | Snapping motion of the jaw of a male that touches a female |
| *"attack male"* | Aggressive | Snapping motion of the jaw of a male that touches another male |
| *“chase female”* | Reproductive | A male moves toward a female and follows the female as it swims away |
| *"chase male"* | Aggressive | A male moves toward a another male and follows the male as it swims away |
| *“dig”* | Reproductive | Fish takes up the gravel in its mouth and expels it outside the territory |
| *“pot entry”* | Reproductive | Fish enters territory (the terracotta pot) |
| *“pot exit”* | Reproductive | Fish exits the territory (the terracotta pot) |
| *“flee from female”* | Aversive | Fish A quickly swims away from a female in direct response to the female |
| *“flee from male”* | Aversive | Fish A quickly swims away from a male in direct response to the female |
| *“lateral display”* | Aggressive | Fish A adopts a lateral orientation, presenting its side to Fish B, often accompanied by fin extension and quivering movements |
| *“lead swim”* | Reproductive | A male fish approaches a female, then reverses direction and guides the female toward the territory. The female follows, with the possibility of entering the territory |
| *“quiver at female”* | Reproductive | A male fish performs rapid, tremulous movements next to a female |
| *“quiver at male”* | Aggressive | A male fish performs rapid, tremulous movements next to another male |

**Supplemental Table 1** Ethogram. List of behaviors, their descriptions, and their assigned behavioral categories

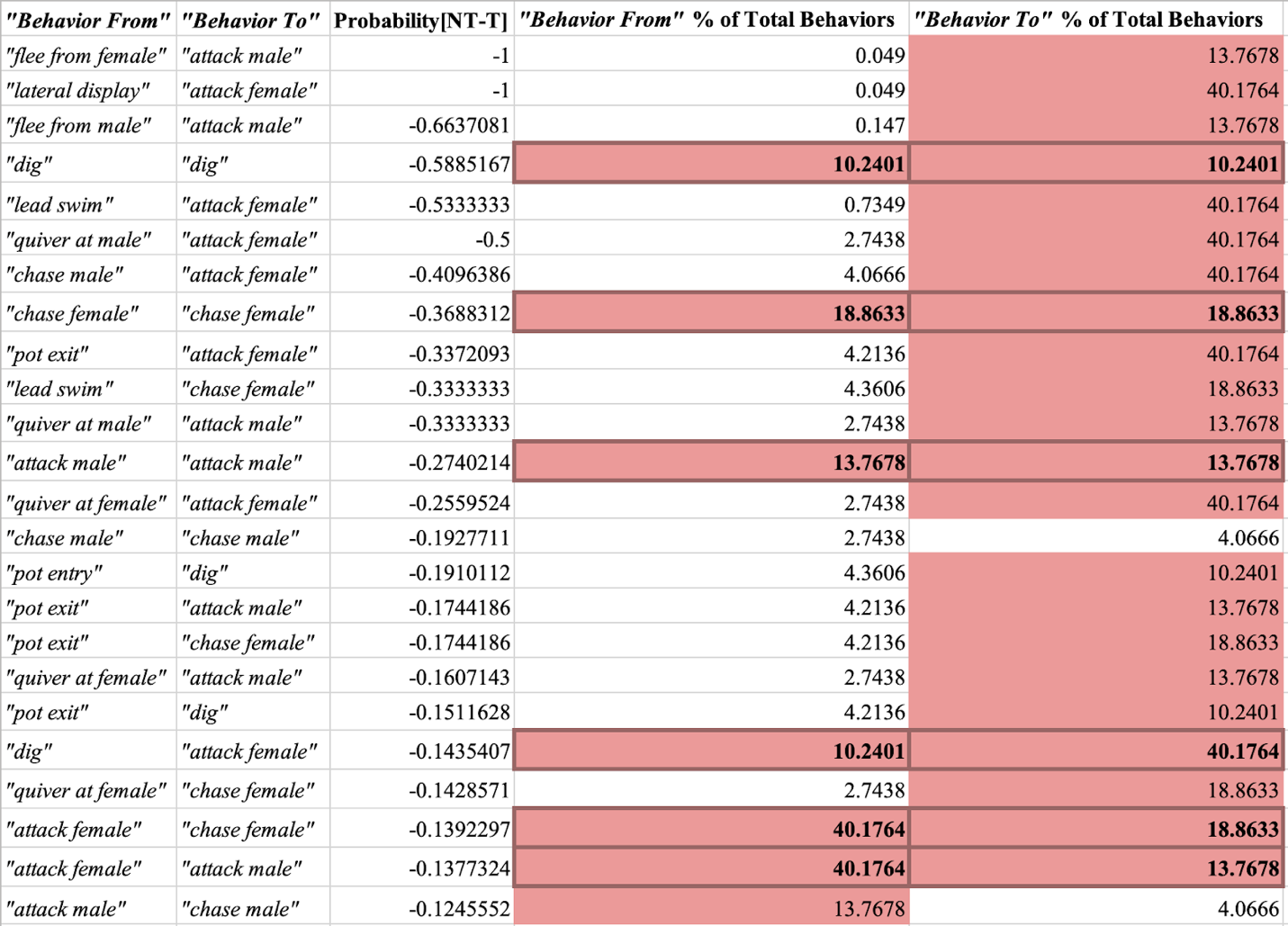

**Supplemental Table 2** List of the behavioral transitions that T males are at least 10% more likely to perform than NTs in the dyad assay. The transitions are separated into their initial and follow-up behaviors, with the incidence of each behavior performed by each subject shown. Behaviors that account for more than 10% of the total behaviors done by Ts are highlighted in red. Dark red squares with bold values indicate that both behaviors in the transition surpassed the 10% incidence threshold. The first column lists the initial behavior, the second column the follow-up behavior, the third column shows the difference in transitional probabilities (p_NT_- p_T_), and the last two columns display incidence of the initial and follow-up behaviors as a percentage, calculated by dividing the relative incidence of each behavior by the total number of behaviors performed by the subject, then multiplying by 100

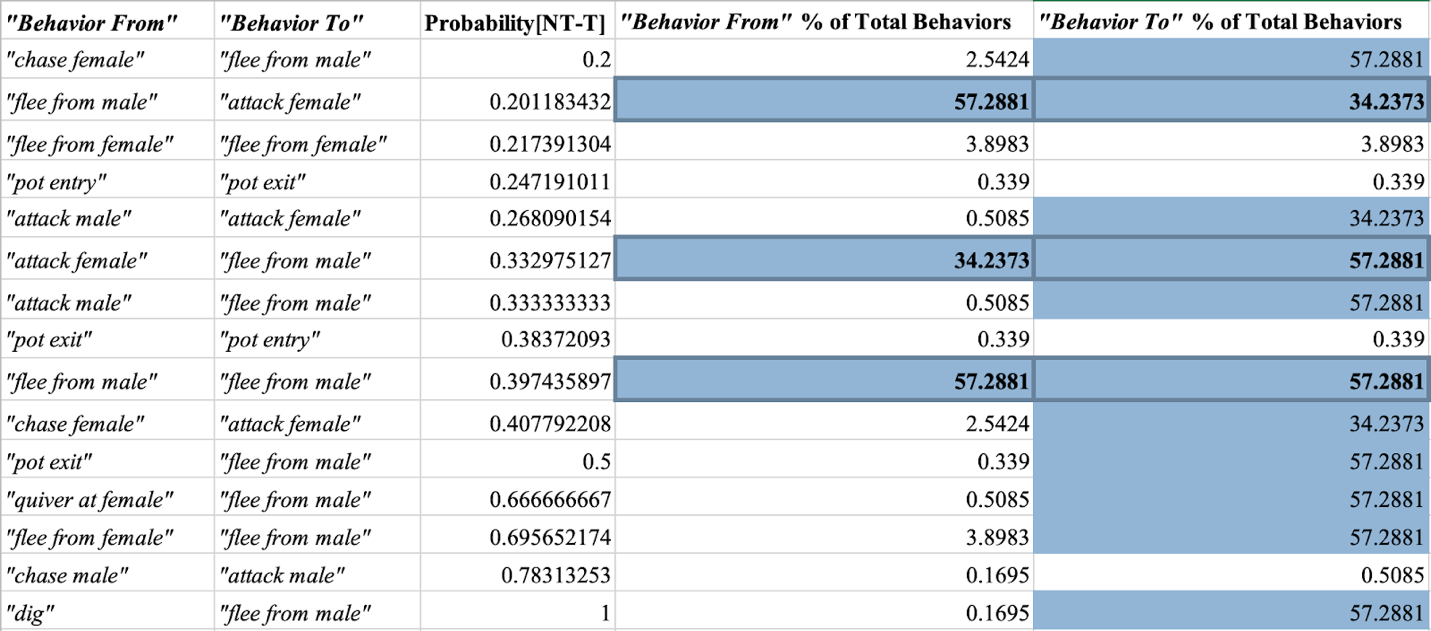

**Supplemental Table 3** List of the behavioral transitions that NT males are at least 10% more likely to perform than Ts in the dyad assay. The transitions are separated into their initial and follow-up behaviors, with the incidence of each behavior performed by each subject shown. Behaviors that account for more than 10% of the total behaviors done by Ts are highlighted in blue. Dark blue squares with bold values indicate that both behaviors in the transition surpassed the 10% incidence threshold. The first column lists the initial behavior, the second column the follow-up behavior, the third column shows the difference in transitional probabilities (p_NT_- p_T_), and the last two columns display incidence of the initial and follow-up behaviors as a percentage, calculated by dividing the relative incidence of each behavior by the total number of behaviors performed by the subject, then multiplying by 100

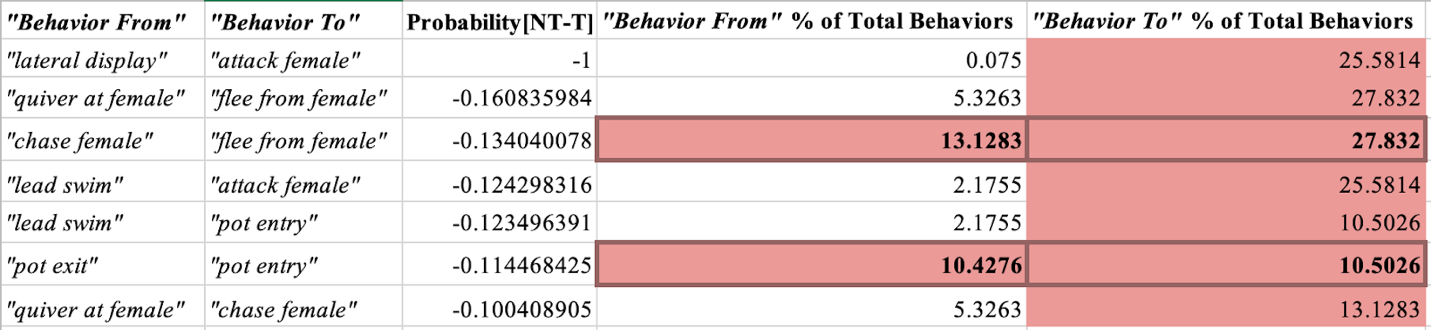

**Supplemental Table 4** List of the behavioral transitions that T males are at least 10% more likely to perform than NTs in the spawn assay. The transitions are separated into their initial and follow-up behaviors, with the incidence of each behavior performed by each subject shown. Behaviors that account for more than 10% of the total behaviors done by Ts are highlighted in red. Dark red squares with bold values indicate that both behaviors in the transition surpassed the 10% incidence threshold. The first column lists the initial behavior, the second column the follow-up behavior, the third column shows the difference in transitional probabilities (p_NT_- p_T_), and the last two columns display incidence of the initial and follow-up behaviors as a percentage, calculated by dividing the relative incidence of each behavior by the total number of behaviors performed by the subject, then multiplying by 100

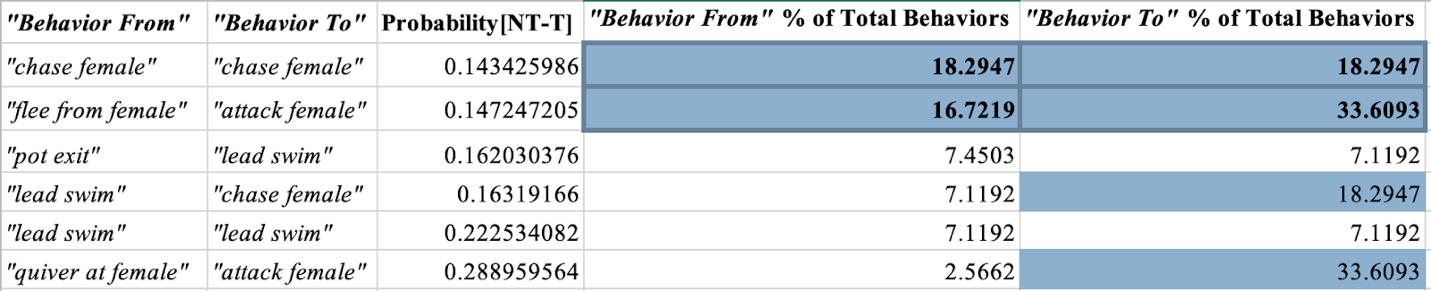

**Supplemental Table 5** List of the behavioral transitions that NT males are at least 10% more likely to perform than Ts in the Spawn assay. The transitions are separated into their initial and follow-up behaviors, with the incidence of each behavior performed by each subject shown. Behaviors that account for more than 10% of the total behaviors done by Ts are highlighted in blue. Dark blue squares with bold values indicate that both behaviors in the transition surpassed the 10% incidence threshold. The first column lists the initial behavior, the second column the follow-up behavior, the third column shows the difference in transitional probabilities (p_NT_- p_T_), and the last two columns display incidence of the initial and follow-up behaviors as a percentage, calculated by dividing the relative incidence of each behavior by the total number of behaviors performed by the subject, then multiplying by 100

| **Behavior From** | **Behavior To** | **Probability T** | **Probability NT** | **Difference** |
| --- | --- | --- | --- | --- |
| *"attack female"* | *"pot exit"* | 0.005865103 | 0 | -0.0059 |
| *"dig"* | *"flee from female"* | 0.015151515 | 0 | -0.0152 |
| *"flee from female"* | *"dig"* | 0.002695418 | 0 | -0.0027 |
| *"flee from female"* | *"pot entry"* | 0.032345013 | 0 | -0.0323 |
| *"lateral display"* | *"attack female"* | 1 | 0 | -1 |
| *"lead swim"* | *"dig"* | 0.034482759 | 0 | -0.0345 |
| *"lead swim"* | *"lateral display"* | 0.034482759 | 0 | -0.0345 |
| *"lead swim"* | *"quiver at female"* | 0.068965517 | 0 | -0.069 |
| *"pot entry"* | *"attack female"* | 0.014285714 | 0 | -0.0143 |
| *"pot exit"* | *"flee from female"* | 0.086330935 | 0 | -0.0863 |
| *"quiver at female"* | *"lead swim"* | 0.014084507 | 0 | -0.0141 |
| *"quiver at female"* | *"pot entry"* | 0.028169014 | 0 | -0.0282 |
| *"dig"* | *"quiver at female"* | 0 | 0.012 | 0.012 |

**Supplemental Table 6** List of behavior transitions that are only occurring within one subject in spawn assay. Behavior transitions highlighted in red, were only performed by the Ts. Transitions highlighted in blue, were only performed by the NTs. “Difference” denotes the difference in probabilities between T and NT [Prob(NT)-Prob(T)]

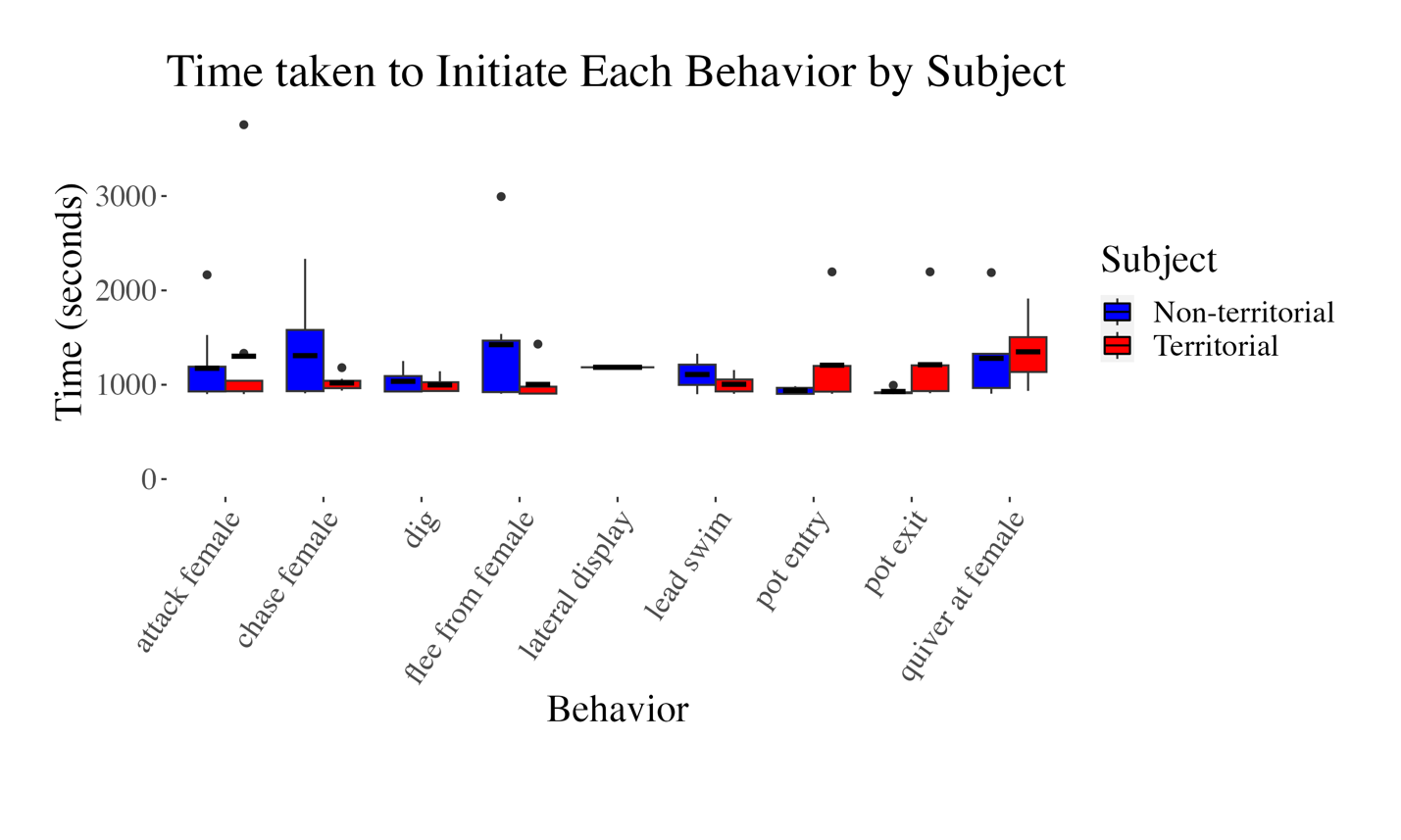

**Supplemental Figure 1** Latency of behavior initiation in spawn assay. Illustration of how quickly Ts and NTs initiated each of the behaviors. There were no significant differences in behavior initiations. Of note, NTs never engaged in lateral display. The box plot illustrates the distribution of data as follows: the upper edge of the box represents the 75th percentile, the horizontal line inside the box marks the mean, and the lower edge of the box represents the 25th percentile. Additionally, the line extending above the box depicts the range of values from the 75th percentile to the maximum, while the line extending below the box shows the range of values from the minimum to the 25th percentile

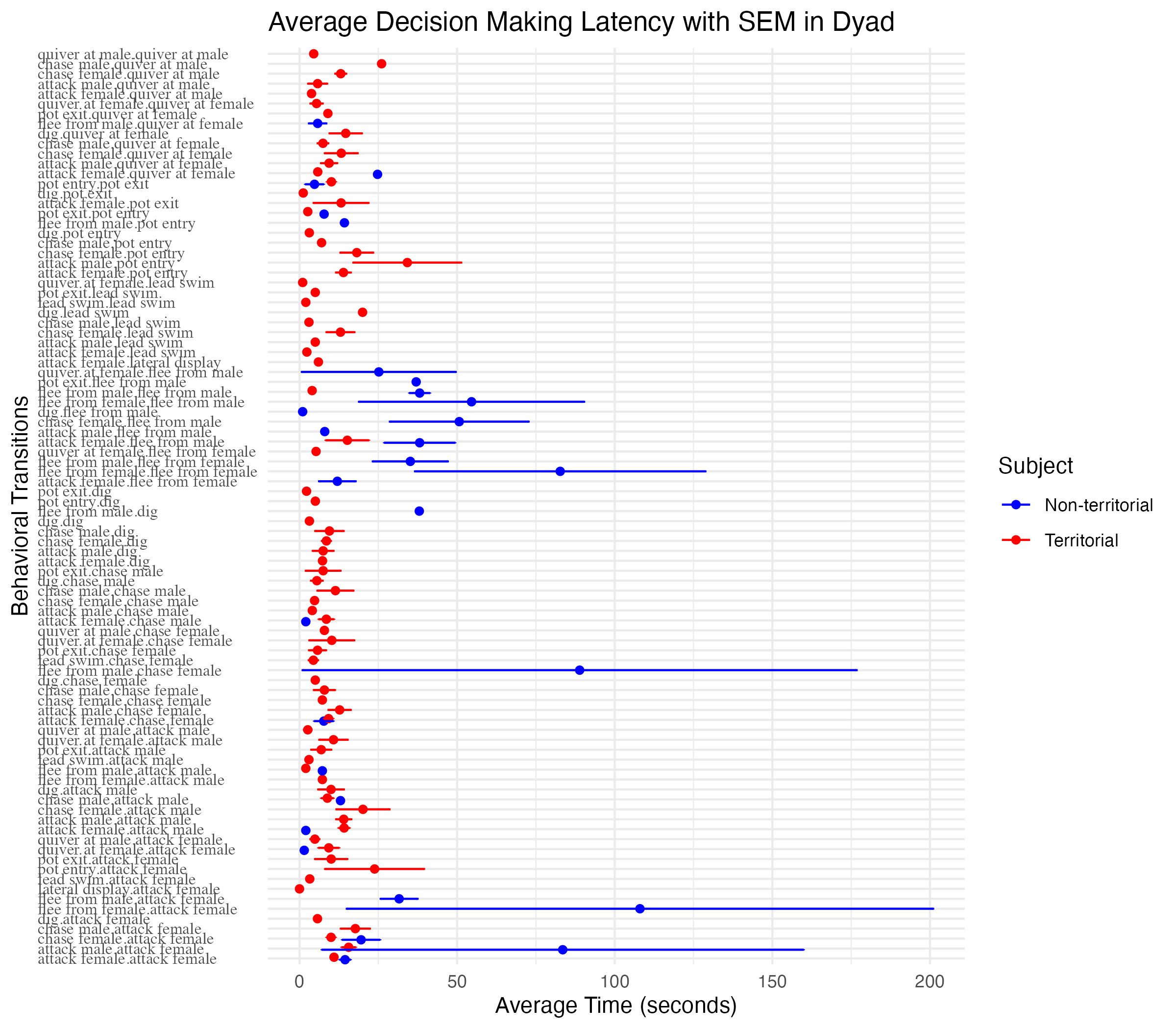

**Supplemental Figure 2a** Decision making latency of T and NT males in dyad assay. This figure illustrates the amount of time, on average, T and NT males took to transition from one behavior to next. Within the behavioral transitions (y-axis), the initial behavior is separated from the subsequent behavior with a period. The red points represent average transition times of territorial (T) males and blue points represent average transition times of non-territorial (NT) males. The lines extending from the points are representing standard error of the mean (SEM)

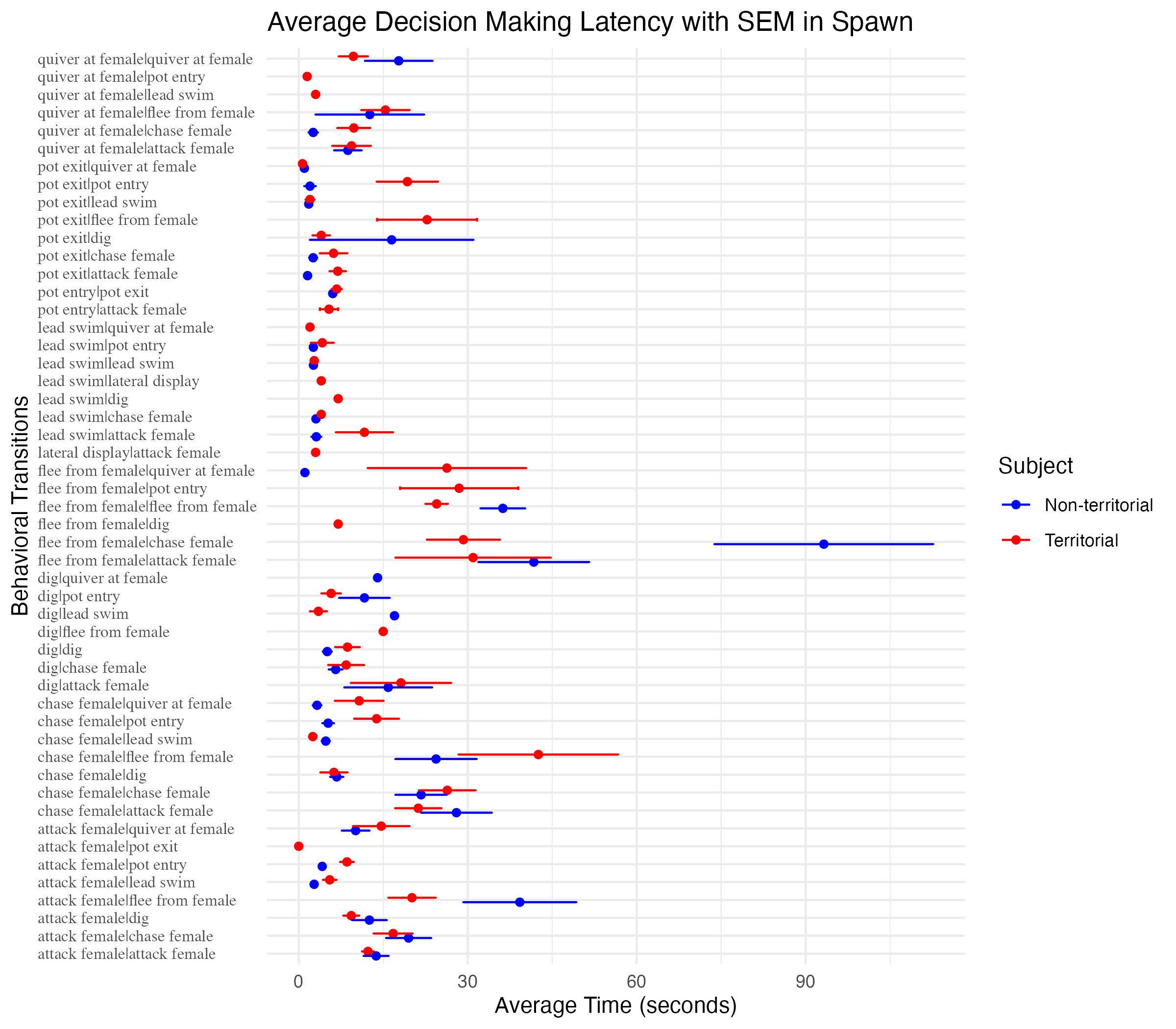

**Supplemental Figure 2b** Decision making latency of T and NT males in spawn assay. This figure illustrates the amount of time, on average, T and NT males took to transition from one behavior to next. Within the behavioral transitions (y-axis), the initial behavior is separated from the subsequent behavior with a period. The red points represent average transition times of territorial (T) males and blue points represent average transition times of non-territorial (NT) males. The lines extending from the points are representing standard error of the mean (SEM)

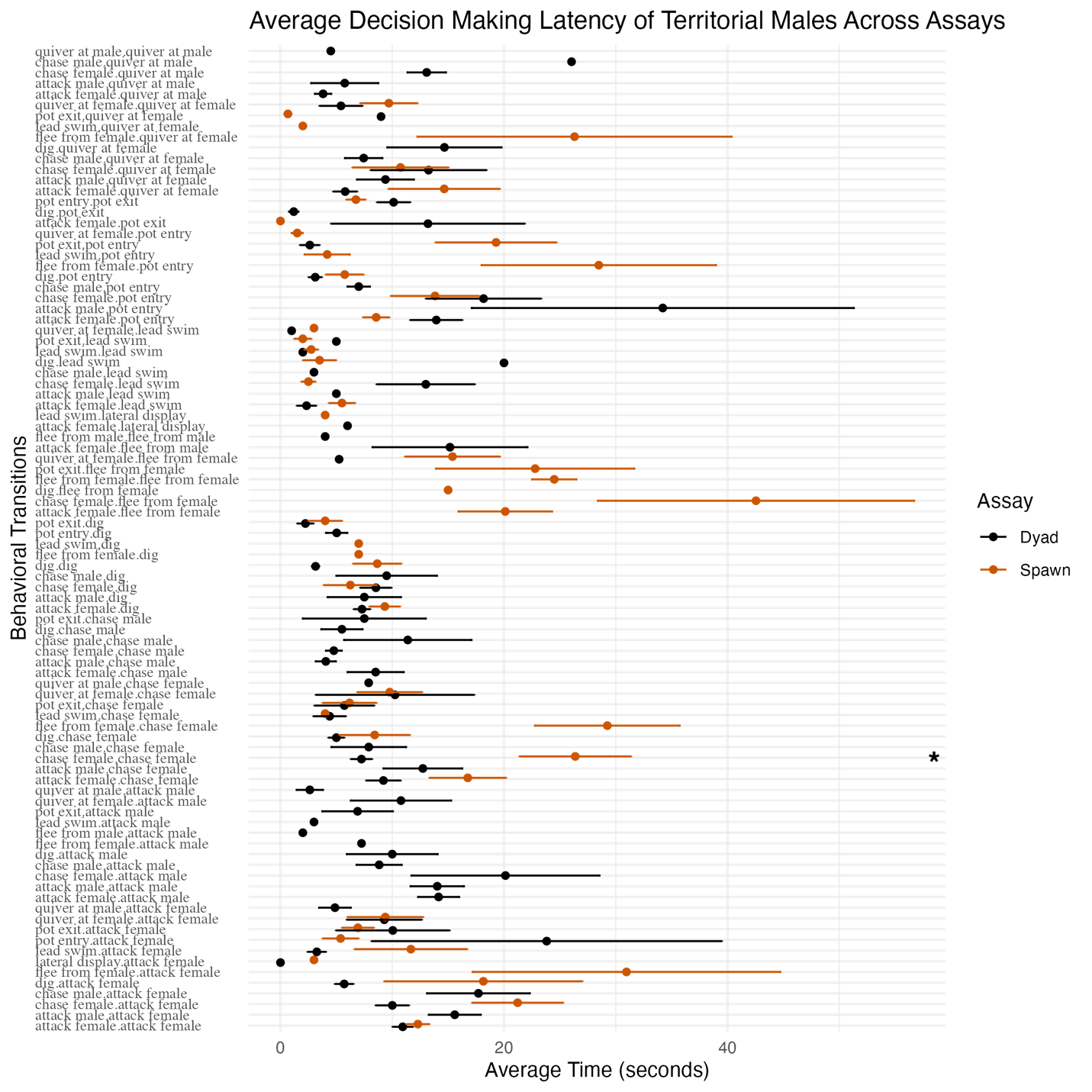

**Supplemental Figure 2c** Decision making latency of T males in dyad and spawn assays. Within the behavioral transitions (y-axis), the initial behavior is separated from the subsequent behavior with a period. The black points represent average transition times of T males in dyad assay and dark orange points represent average transition times of T males in spawn assay. The lines extending from the points are representing standard error of the mean (SEM). Asterisks denote significance (p<0.05)

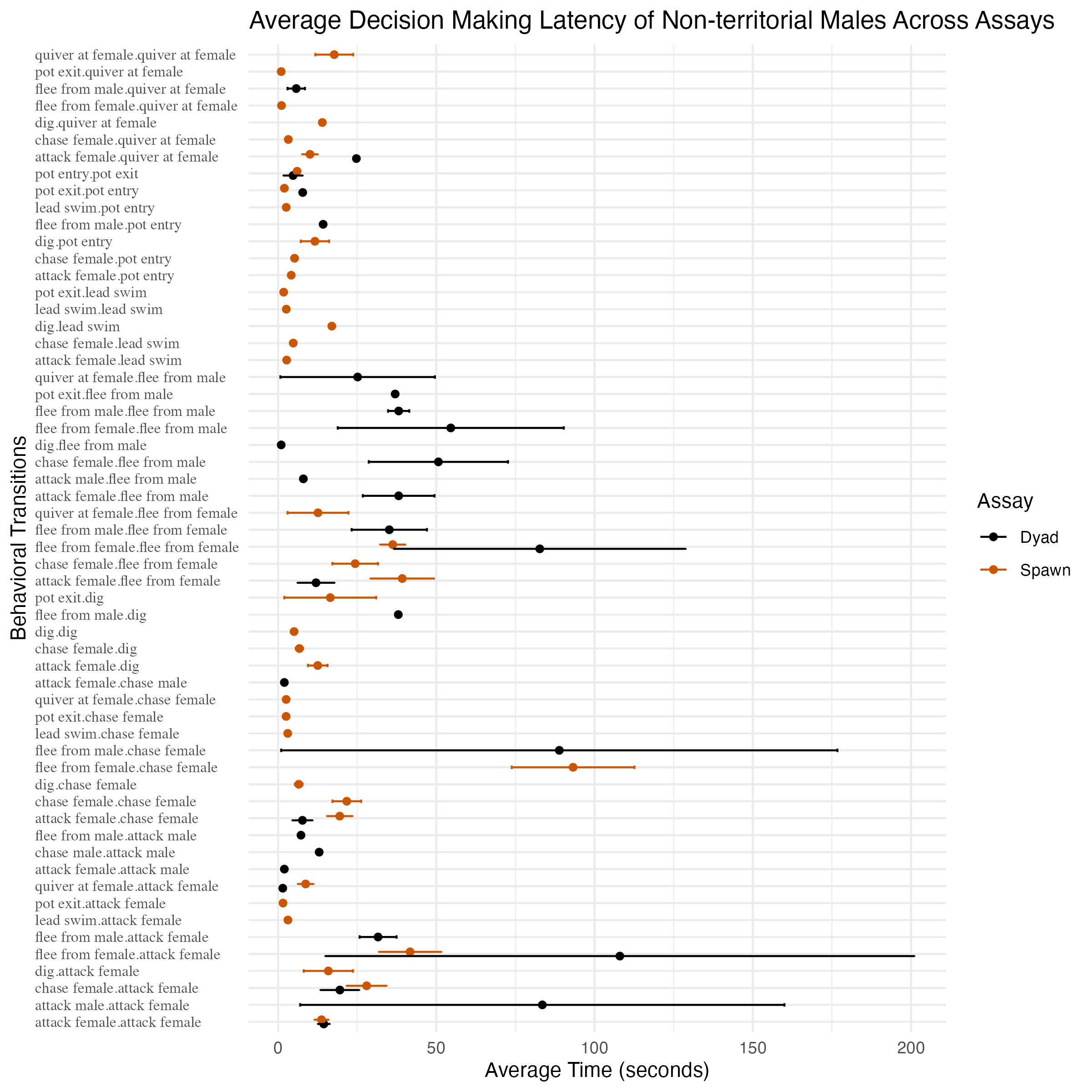

**Supplemental Figure 2d** Decision making latency of NT males in dyad and spawn assays. Within the behavioral transitions (y-axis), the initial behavior is separated from the subsequent behavior with a period. The black points represent average transition times of NT males in dyad assay and dark orange points represent average transition times of NT males in spawn assay. The lines extending from the points are representing standard error of the mean (SEM)

| **Behavior** | **NT Transitional Combinations** | **% of Total Behaviors within previously-NTs** | **T Transitional Combinations** | **% of Total Behaviors within Ts** |
| --- | --- | --- | --- | --- |
| *"attack female"* | 13 | 33.6093 | 16 | 25.5814 |
| *"dig"* | 9 | 6.8709 | 11 | 4.9512 |
| *"pot exit"* | 7 | 7.4503 | 9 | 10.4276 |
| *"chase female"* | 13 | 18.2947 | 13 | 13.1283 |
| *"flee from female"* | 7 | 16.7219 | 11 | 27.832 |
| *"quiver at female"* | 9 | 2.5662 | 11 | 5.3263 |
| *"lead swim"* | 8 | 7.1192 | 12 | 2.1755 |
| *"pot entry"* | 6 | 7.3675 | 9 | 10.5026 |
| *"lateral display"* | 0 | 0 | 2 | 0.075 |

**Supplemental Table 7** Combinatorial centrality in spawn assay. A table describing statistical values that elucidate centrality of T and NT specific behaviors. “Transitional Combinations” refers to how many different combinations each behavior is involved in for T or NT subjects. “% of Total Behaviors” refers to the proportion of each behavior in relation to the total number of incidences for T or previously-NT subjects

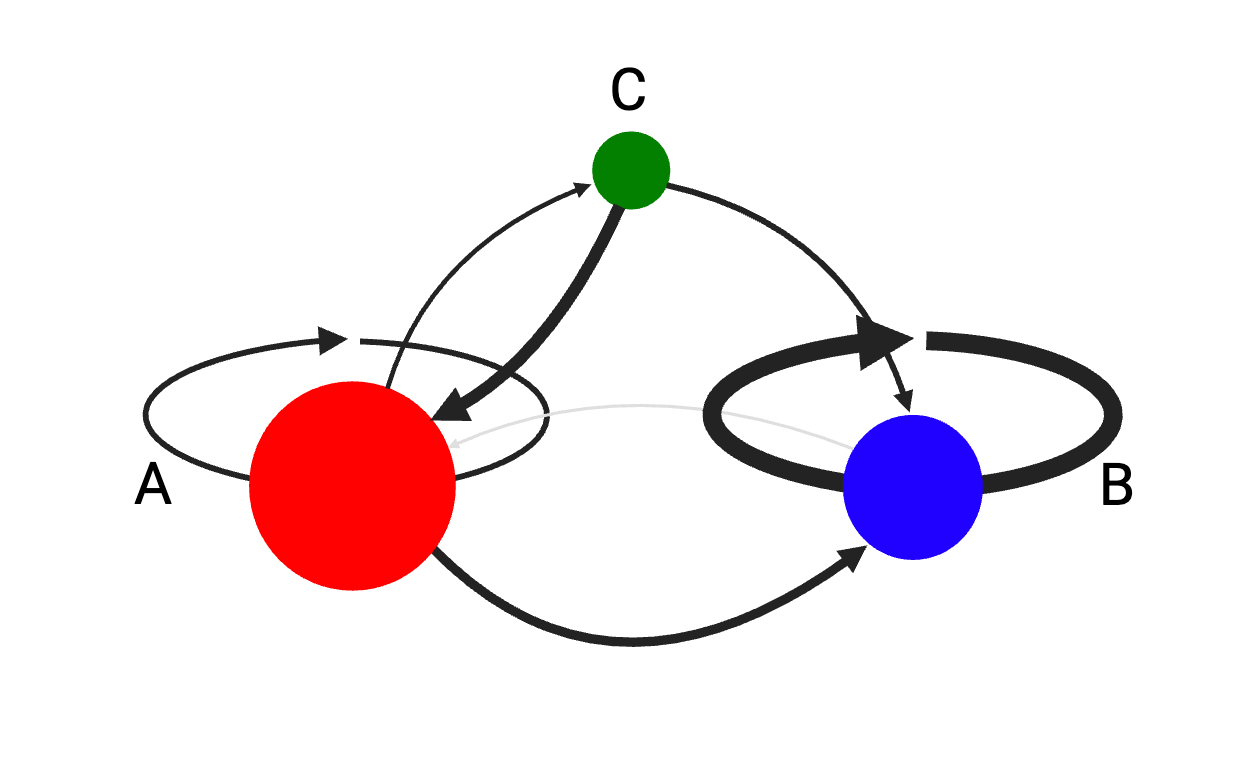

**Supplemental Figure 3** Sample Markov chain figure. Each of the three nodes in the plot correspond to different behaviors. The color of the nodes corresponds to the category of the behavior as specified in the ethogram (see Supplemental Table 1): red for aggressive, green for reproductive and blue for aversive. The relative sizes of the nodes reflect how frequently each behavior occurred. In any given transition, the line originates from the starting behavior and ends at the follow-up behavior with an arrow. Thickness and shade of the connecting lines between nodes represent how likely one behavior is to succeed another: the thicker and darker the line, higher the probability. For example, p_B →B_ is 0.97 while p_B →A_ is 0.03. If a certain node is missing a connecting line to a node, that means the probability of that particular behavioral transition is 0 (e.g. p_C →C_, p_B →C_ )
